# An Explainable Deep Learning Approach for Multimodal Electrophysiology Classification

**DOI:** 10.1101/2021.05.12.443594

**Authors:** Charles A. Ellis, Darwin A. Carbajal, Rongen Zhang, Robyn L. Miller, Vince D. Calhoun, May D. Wang

**Affiliations:** Tri-institutional Center for, Translational Research in, Neuroimaging and Data Science, Georgia State University, Georgia, Institute of Technology, and Emory, University, Atlanta, USA; Wallace H. Coulter Department of, Biomedical Engineering, Georgia Institute of Technology, Atlanta, USA; Department of Computer Information, Systems, Georgia State University, Atlanta, USA; Wallace H. Coulter Department of, Biomedical Engineering, Georgia Institute of Technology and Emory University, Atlanta, USA

**Keywords:** Explainability, Multimodal Fusion, Automated Sleep Staging, Electrophysiology

## Abstract

In recent years, more biomedical studies have begun to use multimodal data to improve model performance. As such, there is a need for improved multimodal explainability methods. Many studies involving multimodal explainability have used ablation approaches. Ablation requires the modification of input data, which may create out-of-distribution samples and may not always offer a correct explanation. We propose using an alternative gradient-based feature attribution approach, called layer-wise relevance propagation (LRP), to help explain multimodal models. To demonstrate the feasibility of the approach, we selected automated sleep stage classification as our use-case and trained a 1-D convolutional neural network (CNN) with electroencephalogram (EEG), electrooculogram (EOG), and electromyogram (EMG) data. We applied LRP to explain the relative importance of each modality to the classification of different sleep stages. Our results showed that across all samples, EEG was most important, followed by EOG, and EMG. For individual sleep stages, EEG and EOG had higher relevance for classifying awake and non-rapid eye movement 1 (NREM1). EOG was most important for classifying REM, and EEG was most relevant for classifying NREM2-NREM3. Also, LRP gave consistent levels of importance to each modality for correctly classified samples across folds and inconsistent levels of importance for incorrectly classified samples. Our results demonstrate the additional insight that gradient-based approaches can provide relative to ablation methods and highlight their feasibility for explaining multimodal electrophysiology classifiers.

## I. Introduction

A growing number of biomedical studies have begun to incorporate data from multiple modalities [1]–[4]. This growth has occurred because complementary modalities increase data richness and can improve classifier performance [2]. However, multimodal data can also increase the difficulty of model interpretation, and many multimodal studies have not incorporated explainability methods that could provide insight into the relative contributions of each modality [3].

A few studies have applied explainability methods to identify the relative importance of each modality to classifiers trained on multimodal data. This includes approaches such as forward feature selection (FFS) [1], impurity [4], and ablation [4][5]. However, some of these methods are incompatible with high-performing deep learning frameworks. FFS requires retraining classifiers repeatedly, so it is not practical for computationally intensive deep learning classifiers. Furthermore, impurity methods can only be applied to decision tree-based models. Unlike FFS and impurity, ablation can be applied to nearly all types of classifiers, is easy to implement, and is not computationally intensive. However, ablation methods do have some weaknesses.

Ablation, like perturbation, requires that the data input to the classifier be modified. This modification can create samples that are out of the data distribution upon which the classifier was trained [6]. Moreover, in deep learning classifiers with automated feature extraction, ablation can cause extracted features that are outside the distribution of other extracted features within the dataset. Furthermore, the goal of ablation is to identify how the performance of a model decreases when the information originally found in a modality is no longer available. As such, it is necessary to be cautious while adapting such methods to new application domains [7]. In domains like electrophysiology (EP) analysis, a value of zero for a modality is abnormal and would likely not bear adequate resemblance to real-life samples. This could affect explainability results, as out-of-distribution or abnormal samples may not correctly assess what a classifier has learned.

Gradient-based feature attribution (GBFA) methods [8] offer an alternative to ablation for multimodal time-series explainability. These methods do not require data modification and are applicable to many deep learning frameworks. Additionally, they can provide much more detailed explanations than ablation. Specifically, ablation can show which modalities were important to the classification of a class, and GBFA methods can show which modalities were important to both the correct and incorrect classification of samples belonging to a class. Layer-wise relevance propagation (LRP) is a popular gradient-based method [9]. Here, we propose the use of gradient-based feature attribution methods, and specifically LRP, for insight into classifiers trained on multimodal data. To demonstrate the viability of this approach, we train a 1-dimensional (1D) convolutional neural network (CNN) for automated sleep stage classification with electroencephalogram (EEG), electrooculogram (EOG), and electromyogram (EMG) data from a popular online dataset [10]. We perform automated sleep stage classification because it is a representative multimodal classification task with clinical needs for model explainability [5]. We apply LRP in a global manner to show the relative importance of each modality to the classification of each sleep stage.

## II. Methods

Here we provide a description of our methods. Using multimodal data, we trained a CNN to discriminate between each sleep stage and applied LRP to explain the decisions of the classifier. The dataset, preprocessing, and classifier that we used here are the same as those which we presented in [7]. The key innovation of this work is its explainability approach.

### A. Description of Data

We used the Sleep Telemetry dataset from the Sleep-EDF Database [10] on Physionet [11]. The dataset included 44, approximately 9-hour recordings from 22 subjects. Each subject had a recording following the administration of a placebo and temazepam. The dataset consisted of EEG, EOG, and EMG with a sampling rate of 100 Hertz (Hz), as well as a polysomnogram of the sleep stage at each time point. For EEG, we used the FPz-Cz electrode. Sleep stages included: awake, movement, rapid eye movement (REM), non-REM 1 (NREM1), NREM2, NREM3, and NREM4. A marker at 1 Hz intervals indicated whether an error occurred in the sleep telemetry device.

### B. Description of Data Preprocessing

We separated each recording into 30-second segments and obtained the corresponding label from the polysomnogram. We discarded movement samples and samples that corresponded with recording errors and consolidated the NREM3 and NREM4 stages into NREM3 [12]. We then z-scored each modality within each recording. The dataset had 42,218 samples and was very imbalanced. Awake, NREM1, NREM2, NREM3, and REM stages composed 9.97%, 8.53%, 46.8%, 14.92%, and 19.78% of the dataset, respectively.

### C. Description of CNN

We adapted a 1D-CNN architecture originally developed for EEG-based sleep stage classification to our multimodal dataset [13]. The architecture, model hyperparameters, and training approach are described in [7]. We used a 10-fold cross validation approach in which training, validation, and test sets were composed of 17, 2, and 3 randomly assigned subjects, respectively. To measure classifier performance, we generated a confusion matrix showing the distribution of sample classification across all folds. Further details on the precision, recall, and F1 score of the classifier are included in [7].

### D. Description of Explainability Approach

We used LRP to explain the relative importance of each modality [9]. LRP provides local explanations for the classification of each individual sample. In LRP, a sample is fed into the neural network and classified. A total relevance of 1 is assigned to the output node for its respective class, and that total relevance is propagated back through the network via LRP relevance rules until a portion of that total relevance is assigned to each of the points in the input sample. Both positive and negative relevance can propagate through the network. Positive relevance shows the features that support the sample being assigned to the class to which it is assigned. Negative relevance identifies the features that support the sample being assigned to other classes. We used the ε and αβ relevance rules [14]. The ε-rule has a parameter, ε, that enables relevance to be filtered when propagated through the network. Increasing ε causes smaller relevance values to be filtered out, reducing the noise in the explanation. The αβ-rule has two parameters, α and β, which control the degree to which positive and negative relevance are propagated through the network, respectively. While the ε-rule allows both negative and positive relevance to propagate, the αβ-rule can enable only positive relevance to be propagated when α equals 1 and β equals 0. We used the ε-rule with an ε of 0.01 and 100 and the αβ-rule with an α of 1 and a β of 0.

To obtain a “global” explanation, we combined the local explanations for all samples in the test set of each fold. We then calculated the percent of absolute relevance assigned to each modality in each fold to identify their relative importance. We did this for all test samples and for each classification group (e.g. awake classified as awake or NREM1 classified as NREM2).

## III. Results and Discussion

Here we describe and discuss the LRP results. We also discuss the study limitations and potential future work.

### A. Explainability Results

Figures 1 and 2 show LRP results for all samples and for each classification group, respectively. When all classes were considered, all LRP rules indicated that EEG was the most important modality, followed by EOG, and EMG. For an ε of 100, when low relevance values were filtered out, the EEG and EOG showed an increase in importance while EMG importance decreased. Interestingly, both the ε-rule (ε=100) and the αβ-rule gave more importance to EMG than the ε-rule (ε=0.01). These results are comparable to the ablation results in [7], though the importance of EOG and EMG appears greater for LRP than for ablation.

**Figure 1.**
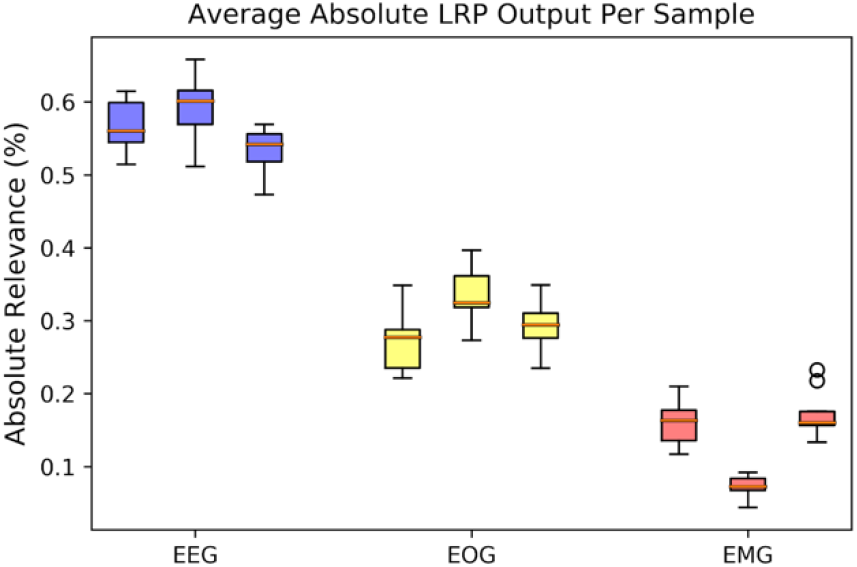
LRP-based global explainability. Plot shows explainability results for all folds. Blue, yellow, and red boxes are for EEG, EOG, and EMG, respectively. Within each trio, from left to right are relevance results for the LRP ε-rule (0.01), ε-rule (100), and α-β-rule.

**Figure 2.**
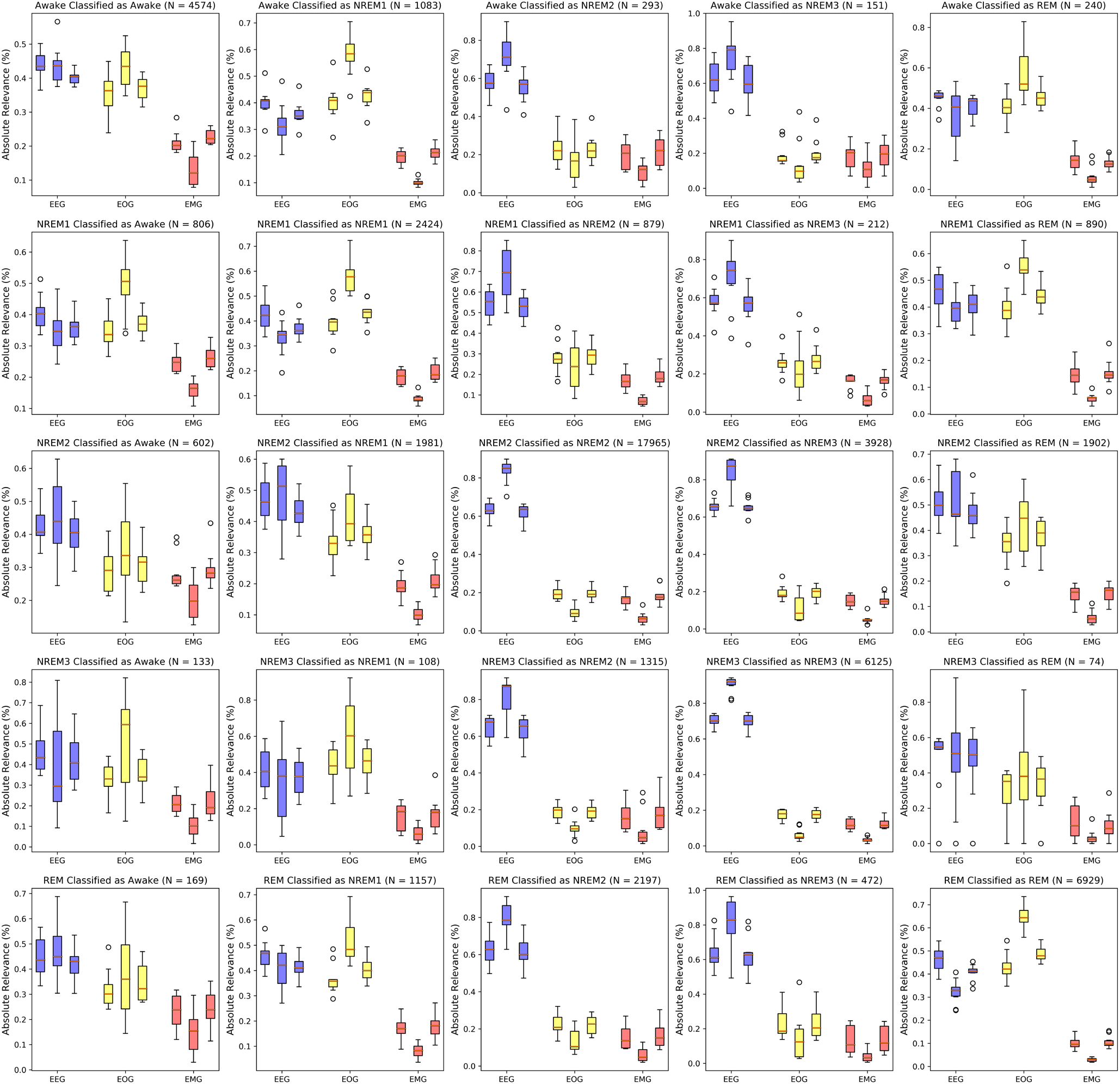
LRP Results for Each Classification Group. Plot shows explainability results for all folds. Blue, yellow, and red boxes are for EEG, EOG, and EMG, respectively. Within each trio, from left to right are relevance results for the LRP ε-rule (0.01), ε-rule (100), and α-β-rule. Note that the number of samples in each classification group is included in the title of each panel.

We also sought to understand the importance of each modality for the correct or incorrect classification of each class. The results in Figure 2 fit with sleep scoring guidelines, as EEG can be used for classifying all stages while EOG and EMG are useful for classifying between awake, REM, and NREM samples [12]. The diagonal of Figure 2 shows the LRP results for correctly classified samples. For the awake stage, the CNN relied mostly on EEG and EOG data. For correctly classifying NREM1, the CNN model placed importance either more on EOG than EEG (ε=100) or about equally on EEG and EOG (ε=0.01 and αβ-rule). However, EMG was the least relevant in correctly classifying NREM1 for all relevance rules (column 2, row 2). For correctly classifying NREM2 (row 3, column 3) and NREM3 (row 4, column 4), EEG had more than 3 times as much relevance as EOG and EMG for all rules. This is consistent with sleep scoring guidelines, as EEG in NREM samples is often very distinct [12]. For correctly classifying REM sleep stages, EOG had the most relevance (ε rule with ε=100 and α-β rule). However, for ε-rule (ε=0.1), EEG and EOG were equally relevant for REM classification. EMG was the least relevant for REM classification across all rules (row 5, column 5). Our results corroborate the well-documented importance of EOG in classifying awake and REM. EOG is important because it tracks eye movements which are more common during awake and REM stages. We also noticed that the relevance across folds of the test samples that the CNN correctly classified had lower variance than the that of the misclassified samples. The lower variance of the correctly classified samples indicates that the features learned by the CNN for correct classification are likely similar across all 10 folds, which could indicate that the architecture is learning generalizable features or that the subjects randomly assigned to each test group are comparable. However, that the CNN seems to have greater variance in relevance across folds for the incorrectly classified samples could indicate that they are making different mistakes in each fold or identifying different ungeneralizable patterns in the training data.

### B. Limitations and Future Work

LRP is one of a broad class of GBFA methods. It is possible that other related methods could provide better explanations. Some metrics have been developed for quantifying the quality of explanations produced by different explainability methods [15], and those metrics could potentially be applied to identify the methods that provide the highest quality explanations for multimodal EP time-series. Furthermore, when we output LRP results, we used each rule for propagating relevance through the whole network. Previous studies have shown that using different rules in different parts of a CNN can improve explanations, especially in deeper networks [14]. Also, we adapted a CNN architecture originally developed for EEG-based sleep stage classification. As such, the architecture was not necessarily developed to extract features optimally from EOG and EMG, which could cause the explainability results to show that EEG is most important. Examining model architectures that might better extract EOG or EMG features could be helpful. Additionally, although our classification performance was below the state of the art, our novel explainability approach, rather than our classifier, was the focus of our study. Using LRP with a better classifier could provide more generalizable explanations and could contribute to novel biomarker identification. Further, local LRP explanations, rather than global explanations, would provide higher resolution insights that might better enable the identification of novel multimodal biomarkers.

## IV. Conclusion

In this study, we implement a gradient-based model-introspection technique for insight into the importance of each modality in multimodal EP data. This offers an alternative to the popular ablation approaches that have previously been used to find the relative importance of each modality to a classifier. Because of its well-characterized clinical guidelines, we used sleep stage classification as a test bed and trained a classifier to discriminate between sleep stages using multimodal data. We further implemented LRP, a popular gradient-based explainability method, to identify the relative importance of each modality to the CNN. Our results corroborate documented findings on the importance of EEG and EOG in classifying awake and NREM1, EOG for REM, and EEG for NREM2-NREM3. They also show that the CNN gave consistent levels of importance to each modality for correctly classified samples across folds and inconsistent importance for incorrectly classified samples. As such, our study demonstrates the additional insight that GBFA methods can provide relative to ablation, highlights their viability for explaining multimodal EP classifiers, and suggests their utility for other multimodal classification problems.

## Acknowledgment

We thank Felipe Giuste and Wenqi Shi for their advice and assistance with computational resources for this project. We thank Mohammad Sendi for helping edit the paper.

